# The transfer function as a tool to reduce morphological models into point-neuron models

**DOI:** 10.64898/2026.03.20.713213

**Authors:** Mikal Daou, Tihana Jovanic, Alain Destexhe

## Abstract

Building a simple model that precisely and functionally characterizes a neuron is a challenging and important task to select the best concise and computationally efficient model. However, this type of work has only been done for subthreshold properties of neurons. Here, we take a different perspective and suggest a method to obtain point-neuron models from morphologically-detailed models with dendrites. To do this, we focus on the functional characterization of the neuron response under *in vivo* conditions, and compute the transfer function of the detailed model. The parameters of this transfer function, in terms of mean voltage, voltage standard deviation and correlation time, can be used to compute the “best” point-neuron model that generates a transfer function very close to that of the morphologically-detailed model. We illustrate this approach for two very different neuronal morphologies, one from *Drosophila* larvae and one from mammals. In conclusion, this approach provides a tool to generate point-neuron models from detailed models, based on a functional characterization of the neuron response.

**Significance Statement:** This study provides a new computational method to reduce morphological models into point-neuron models. To do so, we calculate the transfer function parameters, ie the voltage standard deviation, the mean voltage and the correlation time, of the morphological model and fit a point neuron-model onto this data. Here, we successfully apply this approach for two very different neuron morphologies, a drosophila neuron and a rat motoneuron.

## 1 Introduction

Precise functional characterization of a neuron is a challenging task due to the tradeoff between complexity of the model and performance of the model. Typically, morphological models of neurons can include a lot of details about the neuron, ranging from size, shape and geometry to finer more detailed properties such as channel concentration, types of channels, synapse locations etc. On the other hand, point-neuron models consider the neuron as one homogeneous single compartment. These types of neurons are easier to functionally characterize and to work with. Finding a method that allows us to use a point-neuron model that incorporates computational properties identified in a morphological model can considerably improve the precision of single neuron models. For example, we could incorporate precise quantal or density information about synapses in our models.

A way to functionally and efficiently characterize a neuron is to build a simple model that can predict its firing rate based on the frequencies of excitatory and inhibitory inputs. The transfer function defined in Zerlaut et al. (2016); El Boustani and Destexhe (2009) has been shown to do that very well for adaptive exponential integrate and fire models (Brette and Gerstner, 2005). However, this type of characterization has never been applied to detailed morphological models. The single-compartment approximation enables analytical tractability and efficient numerical computation.

Here, we use a novel approach to compute the transfer function of a morphologically-detailed model and find the best corresponding point-neuron model (figure 1). This allows us to consider how the input-output relationship is affected by morphological properties such as geometry, variation of channel concentration and passive properties along the neuron, synapse locations, dendritic spikes, etc. We can focus this question on specific neuron types or neurons from specific species and compare the resulting transfer functions across these different morphologies. That can be interesting because some neurons have unique morphological properties. For instance, active channels in dendrites have been frequently reported only in mammalian neurons where they have been shown to have a great electrophysiological and computational role (Magee and Johnston, 1995; Stuart et al., 1997; Markram et al., 1995). On the other hand, insect neurons, for example, have distinct morphological properties that differ greatly from mammalian neurons such as the distance between the soma and the axon relative to the size of the neuron (Cardona et al., 2010; Jovanic et al., 2019). One striking property of insect neurons is the fact that the axoemerges from a common primary process called neurite while the soma is located at the other end of the neuron (or neurite) (Ananthakrishnan, 1976). There have been very little electrophysiological studies of insect neurons, so the computational consequences of these properties remain unknown.

**Figure 1:**
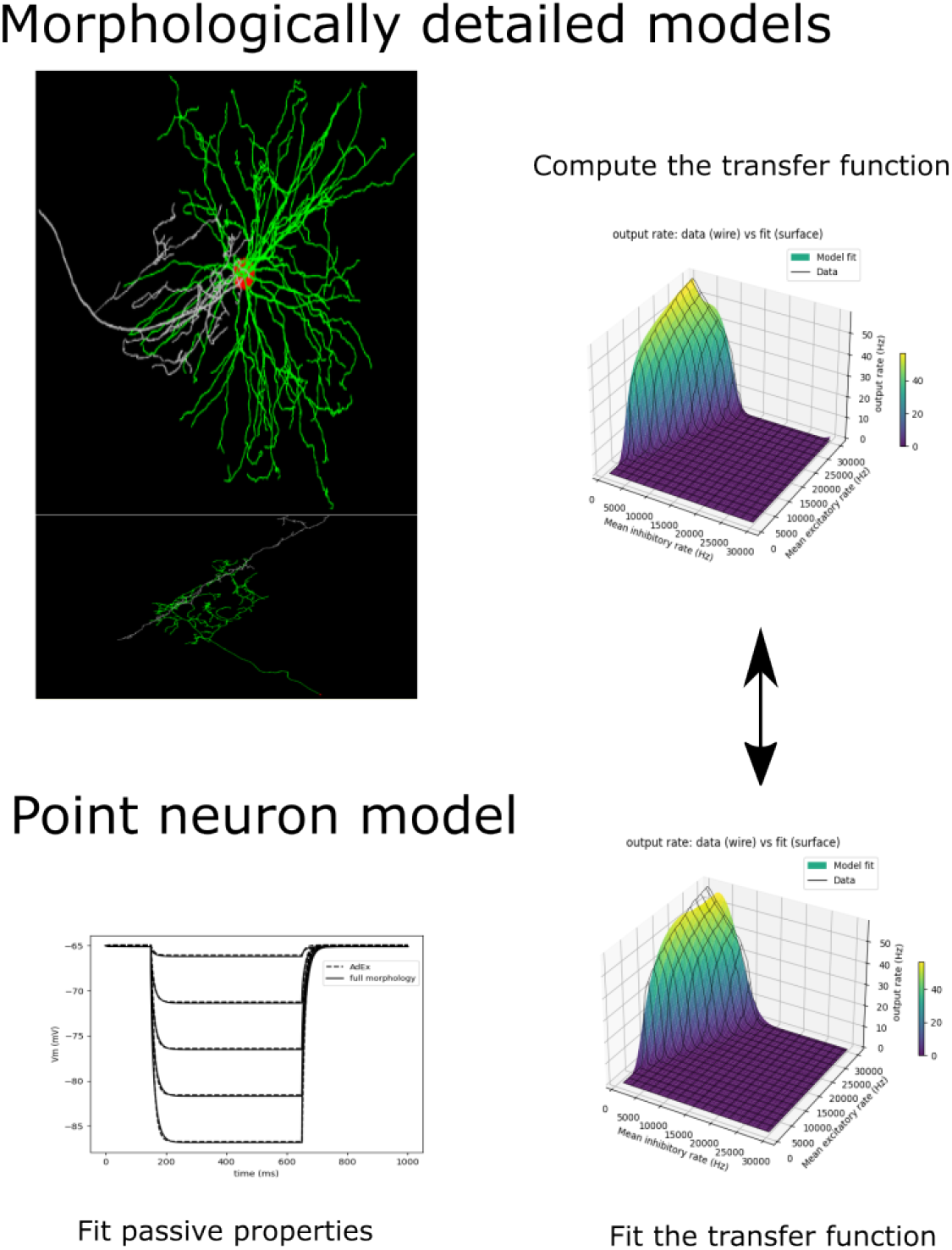
transfer function and point-neuron fitting pipeline. The transfer function is first computed from the morphological model (top). The point-neuron model (bottom) is first fit to the detailed model for the passive properties (bottom left), then the transfer function is calculated and fit to that of the detailed model.

Furthermore, we wanted to see if the transfer function of a morphological model can be used as a proxy to find the best matching point-neuron model. Finding the best point-neuron model that describes a detailed morphological model is a challenging task that often requires methods of model reduction (Di Volo et al., 2019) which can lead to a loss of information. Fitting the transfer function of a point-neuron model to statistics of a morphological model can be a way to accurately transition from a detailed model to a simplified one while guaranteeing that both are functionally equivalent

Here, we propose a method to collapse the original morphological model into a point-neuron model with the same input-output transfer function. As schematized in Fig. 1, we first calculate the transfer function of the morphologically detailed neuron under *in vivo* conditions. The simplified point-neuron model is then first adjusted to fit the passive properties of the detailed neuron, and in a next step, we calculate its transfer function by varying the synaptic parameters, and finally fit to the detailed neuron’s transfer function. We show the application of this method for two different types of neurons, one from a mammal and one from a *Drosophila*.

## 2 Methods

### 2.1 Parameter choice

All simulations and fitting procedures were run in NEURON by varying membrane resistance, axial resistance, and capacitance. The parameters were assumed to be uniform across the cells. For Basin-2, the fitting was done using NEURON’s fitting algorithm PRAXIS. For the motoneuron, we randomly assigned some parameter values from a biologically plausible range based on available data.

### 2.2 Calculation of the transfer function

The transfer function is defined in Zerlaut et al. (2016) as:

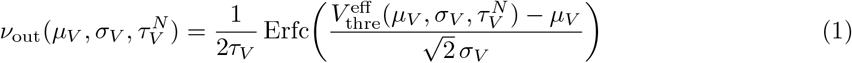

where Vout is the firing rate and the parameters (*µ*_*v*_, *σ*_*v*_, *τ*_*v*_) are functions of (*ν*_*e*_, *ν*_*i*_, *w*). *µ*_*v*_ is the mean voltage, *σ*_*v*_ is the standard deviation, *τ*_*v*_ is the autocorrelation time constant, *ν*_*e*_ is the mean excitatory input frequency, *ν*_*i*_ is the mean inhibitory input frequency, and *w* is the adaptation current of the transfer function. *µ*_*v*_, *σ*_*v*_, *τ*_*v*_ were calculated through simulations by varying the inhibitory *ν*_*i*_ and excitatory *ν*_*e*_ input frequencies from 1 to 30 kHz. For each combination of *ν*_*e*_ and *ν*_*i*_, we ran a 2 second simulation at the end of which we calculated the parameters.

In Zerlaut et al. (2016), the effective threshold 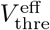 has been defined as a second order polynomial as follows:

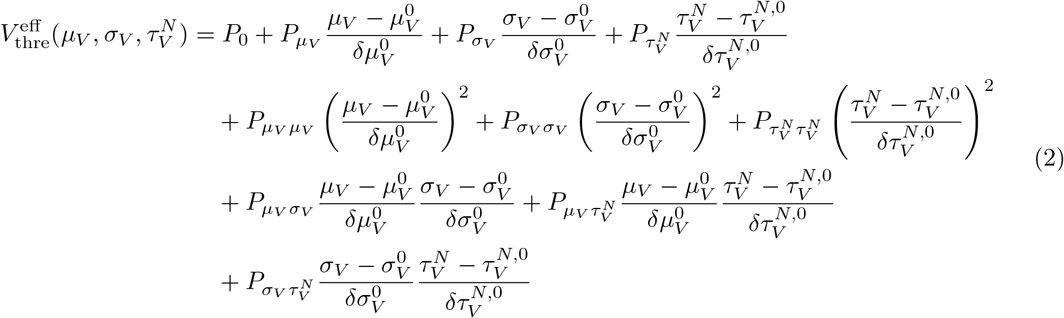

The values used here for the polynomial parameters are those used in Di Volo et al. (2019). We then calculate P by fitting Vout to the firing rates calculated through the simulations.

### 2.3 Fitting a point-neuron model

Using the AdEx model, we then tried to find the best point-neuron model that has the same transfer function. We simulated step current injections in each neuron to select to fit *C*_*m*_, *E*_*L*_ and *g*_*l*_. We then fitted the theoretical distributions of *µ*_*v*_, *σ*_*v*_, *τ*_*v*_ on the data calculated through simulations of the morphological model with *Q*_*e*_, *Q*_*i*_, *τ*_*e*_, *τ*_*i*_ being the free parameters. We then used the all the fitted AdEx parameters to recompute the threshold polynomial using the semi-analytical method and we compared the resulting firing rate distribution with the first one.

### 2.4 Synapse distribution

For Basin-2, we used data from electron microscopy reconstruction of the larval CNS (central nervous system) performed in Catmaid (Ohyama et al., 2015; Saalfeld et al., 2009) to distribute synapses and weights coming only from a specific sensorimotor network in which the neuron is involved (Jovanic et al., 2016). That allows us to study the neuron’s firing properties in this specific network. For the rat motoneuron, we defined specific regions to which we assigned a value for synaptic density based on the synapse type (excitatory vs inhibitory) and using realistic biological ranges. The 3 defined regions are the soma, the proximal and dendrites.

## 3 Results

### 3.1 Calculation of the transfer function a morphological model of Basin-2

There were very few electrophysiological studies of *Drosophila* neurons. We therefore chose Basin-2, an interneuron neuron in a sensorimotor network in the *Drosophila* larval ventral nerve cord (Ohyama et al., 2015; Jovanic et al., 2016). That would indeed constitute one of the first functional characterizations of a *Drosophila* neuron and that could be used to compare these neurons with more widely studied neurons from mammals. The first step for computing the transfer function of the 3d model of this neuron is to calculate the parameters of the transfer function (*µ*_*v*_, *σ*_*v*_, *τ*_*v*_) and the firing rates for each combination of (*ν*_*e*_, *ν*_*i*_) through simulations (figure 2). We can then fit the polynomial function (2) used to calculate the effective threshold in (1) and recalculate the firing rates semi-analytically. Based on electron microscopy reconstruction data from Catmaid, we only placed synapses from a network involved in sensori-motor control (Jovanic et al., 2016) to see if we could specifically study the firing statistics of this neuron involved only in one network. We managed to obtain a good fit for all three parameters of the transfer function as well as for the firing rates (figures Figures 2,3).

**Figure 2:**
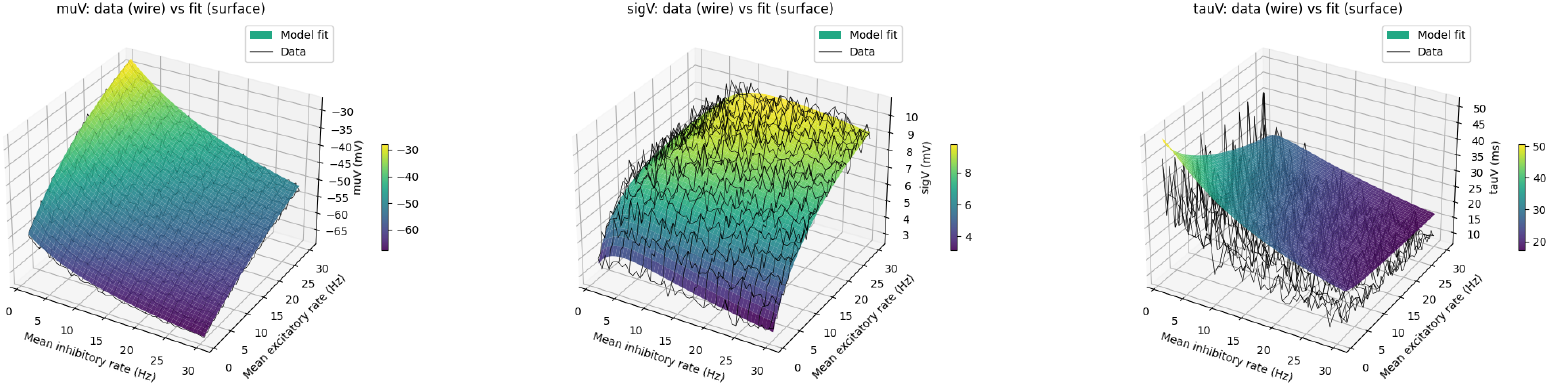
3d plots of the transfer function parameters fit for Basin-2. Computation of the voltage moments through simulations (black surfaces) and using the full transfer function pipeline (colored surfaces) after having fitted the AdEx parameters. *µ*_*v*_ (left), *σ*_*v*_ (middle), *τ*_*v*_ (right) are computed as a function of inhibitory and excitatory synaptic rates.

**Figure 3:**
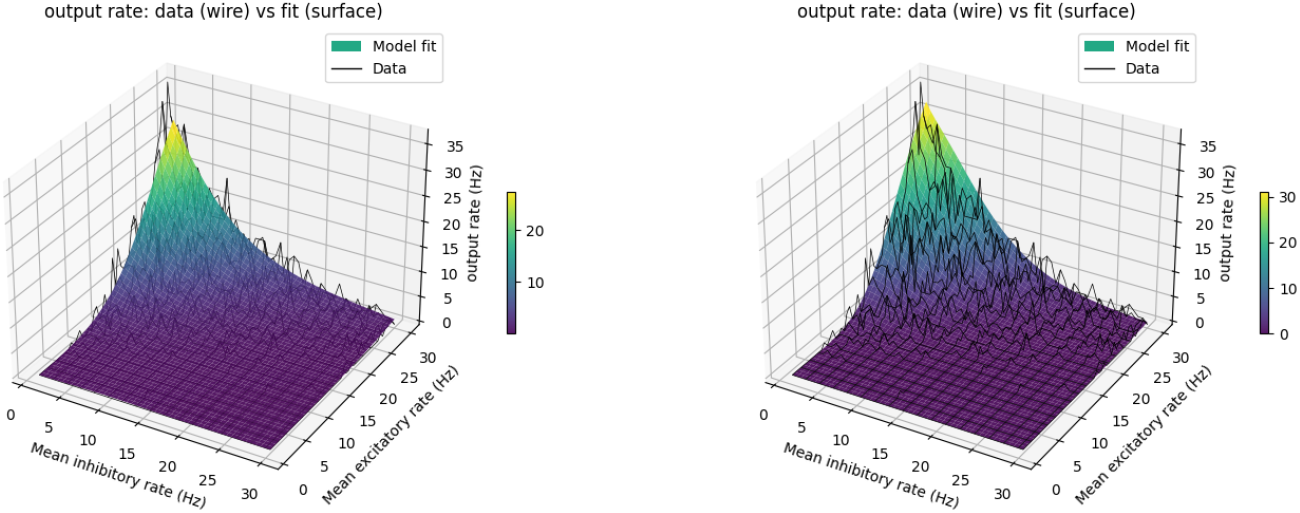
3d plots of the observed (black surfaces) and predicted (colored surfaces) firing rates for Basin-2. Firing rates are first computed through simulations as a function of inhibitory and excitatory synaptic rates. The transfer function polynomial is then fitted to calculate the firing rates using the parameter calculated through simulations (left) and using the full transfer function pipeline after having fitted the AdEx parameters (right). The same polynomial was used for both cases.

We then wanted to see if the transfer function could be used to find a point-neuron model that best reproduces the firing properties of the morphological model. To do so we used the transfer function equations that are used for the AdEx model to fit an AdEx point-neuron on the statistics that were computed through simulations. We first fitted the passive properties of the AdEx onto the morphological model by simulating a current clamp procedure (figure 4). We then fixed the synaptic time constants to 5 ms and varied *Q*_*e*_ and *Q*_*i*_ to fit *µ*_*v*_, *σ*_*v*_ and *τ*_*v*_ calculated from the AdEx model onto the same statistics calculated through simulations of the morphological model. The match between the data and the theoretical distributions was good enough that we could use the same polynomial to compute the transfer function from the AdEx (figure 3).

**Figure 4:**
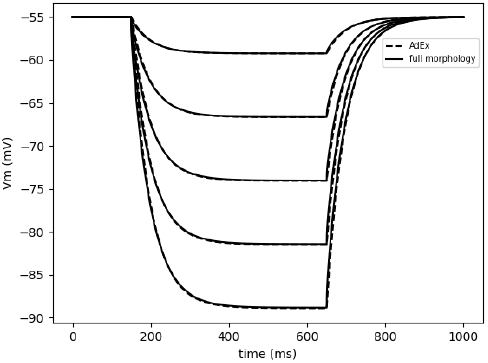
Current clamp simulations in the AdEx model and the morphological model of Basin-2. Current clamp injections were run for the morphological model (filled curves) and the AdEx model (dashed curves) to fit the passive parameters of the AdEx model. Current injections range from -0.016 nA to -0.002 nA.

### 3.2 Calculation of the transfer function of a rat motoneuron

We also wanted to see how our results vary across various morphologies and synapse distributions, so we then calculated the transfer function of a rat motoneuron for various synapse distributions. To do so, we divided the morphology into 3 regions and assigned a synaptic density value for each of them. We used the same method as with Basin-2 to fit an AdEx model by first fitting the transfer function using simulation statistics (figure 5), then fitting the passive properties (figure 6) and calculating the firing rates semi-analytically (figure 7). However, the slight discrepancies between the voltage moments calculated analytically and those calculated through simulations forced us to calculate the polynomial again when fitting the transfer function on the AdEx model.

**Figure 5:**
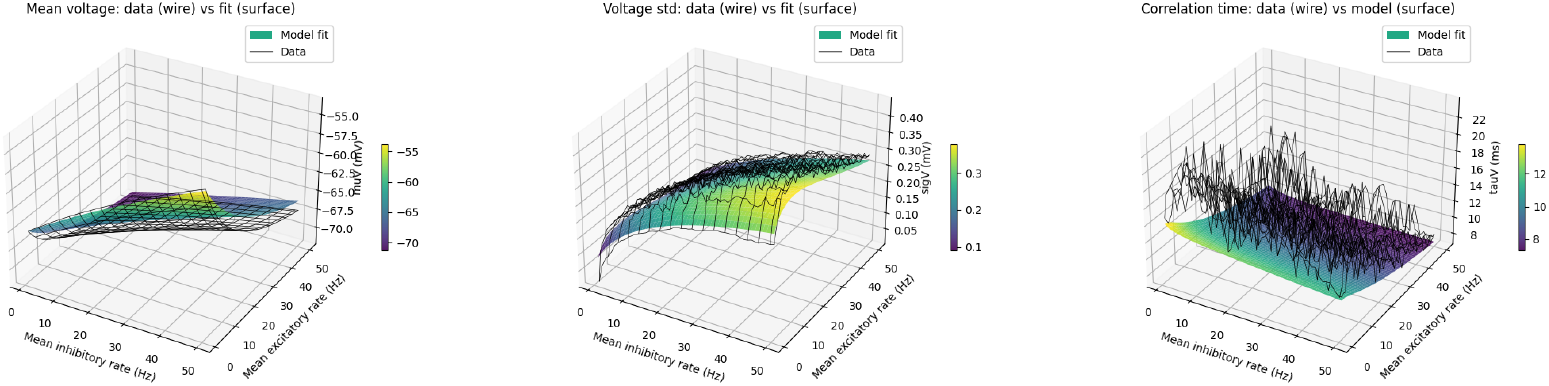
3d plots of the transfer function parameters fit for a rat motoneuron. Computation of the voltage moments through simulations (black surfaces) and using the full transfer function pipeline (colored surfaces) after having fitted the AdEx parameters. *µ*_*v*_ (left), *σ*_*v*_ (middle), *τ*_*v*_ (right) are computed as a function of inhibitory and excitatory synaptic rates.

**Figure 6:**
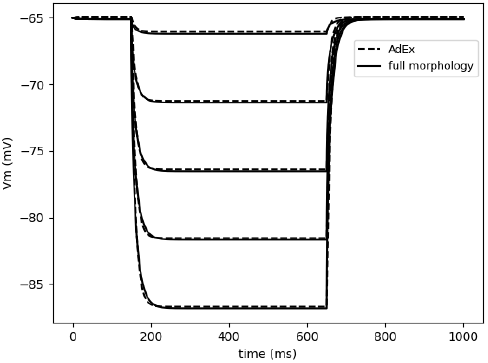
Current clamp simulations of the morphological model and the point-neuron model of a rat motoneuron. Current clamp injections were run for the morphological model (filled curves) and the AdEx model (dashed curves) to fit the passive parameters of the AdEx model. Current injections range from -2 nA to -0.1 nA.

**Figure 7:**
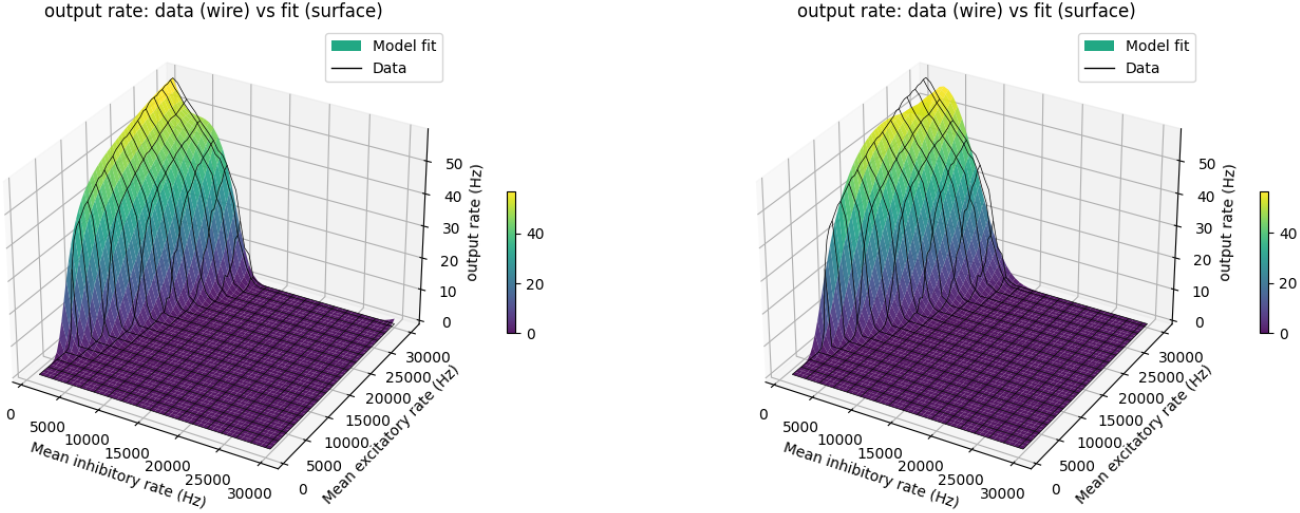
3d plots of the transfer function fit for a rat motoneuron. Firing rates are first computed through simulations as a function of inhibitory and excitatory synaptic rates. The transfer function polynome is then fitted to calculate the firing rates using the parameter calculated through simulations (left) and using the full transfer function pipeline after having fitted the AdEx parameters and having fitted the polynomial again(right).

To evaluate the match between the original morphological model and the fitted AdEx model, we compared simulations of the two models for the same synaptic Poisson input frequencies. We used the passive parameter values that were fitted during the current clamp procedure and the synaptic conductances and time constants optimized when fitting *µ*_*v*_, *σ*_*v*_ and *τ*_*v*_. However, given that the *V*_*T*_ in the transfer function formalism is dynamic and defined as a function of (*ν*_*e*_, *ν*_*i*_), we varied *V*_*T*_ and ran simulations for each combination of Poisson synaptic and inhibitory rates to compute the firing rate distribution and compare it with the original one (figure 9). we then compared actual simulations of both models for specific combinations of inputs (figures 8) and found similar firing frequencies between both models.

**Figure 8:**
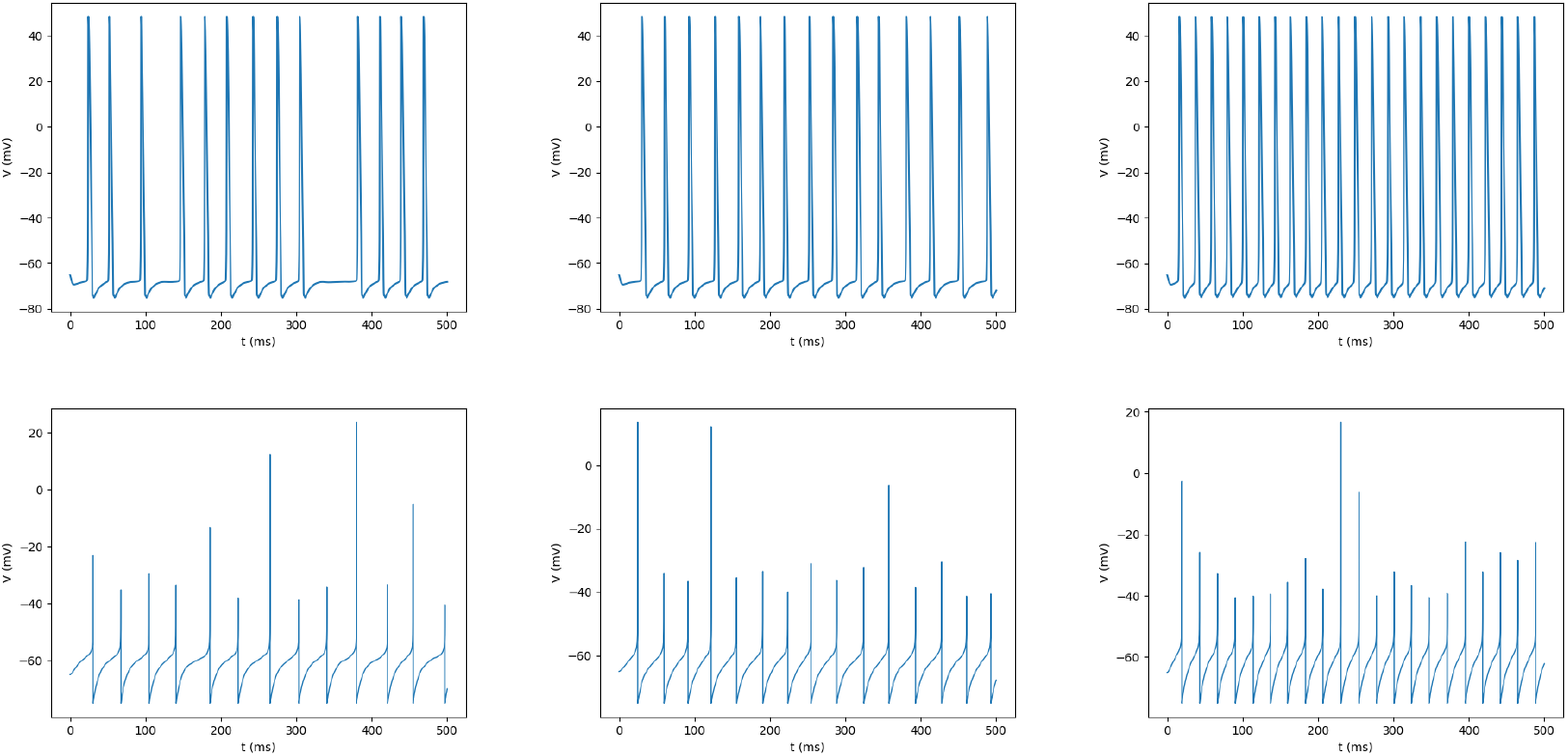
Comparison of simulations of the morphological model with simulations of the AdEx model. synaptic input frequencies from left to right: *ν*_*e*_=30 Khz and *ν*_*i*_=6.3 Khz, *ν*_*e*_=15.8 Khz and *ν*_*i*_=1 Khz, *ν*_*e*_=30 Khz and *ν*_*i*_=1 Khz. Firing frequencies for the morphological model (top) from left to right are: *ν*_*out*_=13 Hz, *ν*_*out*_=30 Hz, *ν*_*out*_=46 Hz. Firing frequencies for the AdEx model (bottom) from left to right are: *ν*_*out*_=26 Hz, *ν*_*out*_=30 Hz, *ν*_*out*_=42 Hz

**Figure 9:**
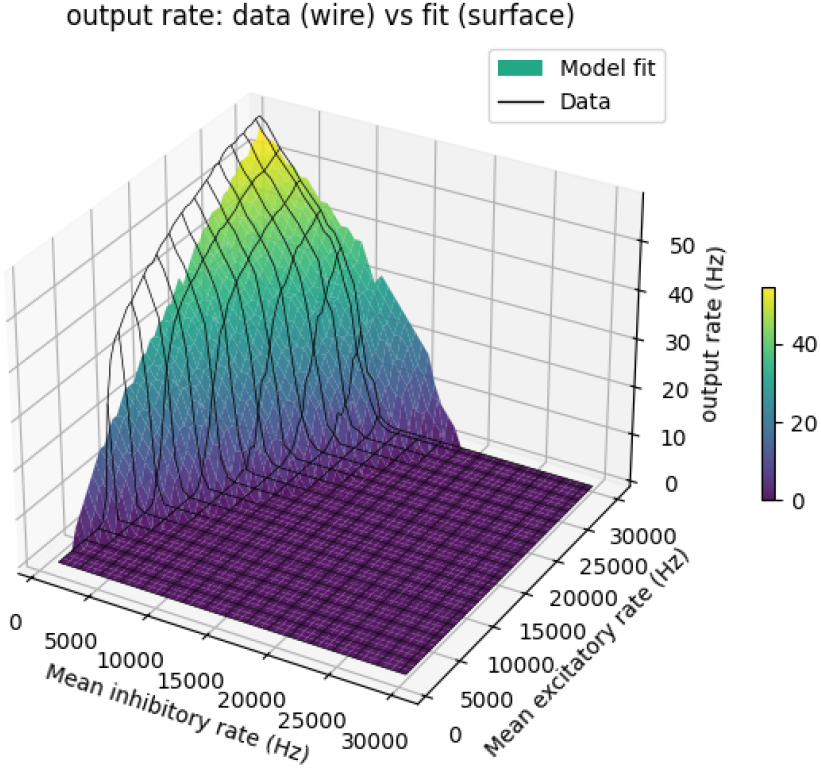
Plot of the firing rates of the AdEx model calculated through simulations (black surface) and original firing rates (colored surface). Here, the firing rates of the AdEx model were calculated by running a simulation for each combination of synaptic input frequencies and calculating *ν*_*out*_, just like originally done with the morphological model.

## 4 Discussion

In this paper, we developed a novel approach that allows us to functionally characterize morphological models and find the best matching point-neuron model. We do so by calculating the transfer function of the morphological model and finding the best AdEx model that has the same transfer function.

### 4.1 Comparison to previous approaches

The first method developed to design simplified models of detailed morphological models is Rall’s equivalent cable, which collapses the neuron’s dendritic tree into a single cable (Rall, 1959). However, the morphology needs to satisfy some constraints for the method to be applicable: each branch points must satisfy the 3/2 power law, the membrane is purely passive and uniform, the membrane potential is assumed to be uniform and all dendrites must have the same length to the soma. The resulting model obtained by this method, consists of a soma coupled to a single cylinder for each dendritic tree, and not one single compartmental model.

A second method is compartmental pruning (Bush and Sejnowski, 1993) which iteratively removes compartments while adjusting parameters to end up with a reduced model that preserves the electrical properties of the original model. This method is purely empirical, computationally expensive, still doesn’t lead to a single compartmental model and the result depends on the pruning order.

A third type of method is based on preserving dendrite-to-soma attenuation (Destexhe, 2001; Wybo et al., 2021). In such methods, the reduced model is constrained by capturing the attenuation of responses along the dendrites and obtaining correct somatic responses in the reduced model. This can be done by preserving membrane area, input resistance, time constant, passive responses, voltage attenuation and somatic responses (Destexhe, 2001). Similar methods use the soma-dendrite impedance function of the neuron to constrain the simplified model. For instance, in Wybo et al. (2021); Yan and Li (2011), the method preserves the input-output behavior by treating the neuron as a linear dynamical system around a subthreshold operating point. The resulting model is a reduced RC-network or 2-5 compartments. These methods preserve the global electrical properties such as input resistance and membrane time constants but the compartments have no anatomical meaning.

Active reduced methods (Pinsky and Rinzel, 1994; Mainen and Sejnowski, 1996; Bush and Sejnowski, 1993) go a bit further by collapsing the original model into a simpler one that includes a few key morphological compartments such as the soma and a few dendrites that are functionally distinct from each other. In Rössert et al. (2017), the passive properties of the point neuron, spiking behavior and synaptic properties are fit on current injection simulations of the morphological model. However, the identified single compartmental model is fitted in a restrictive regime the voltage moments such as the mean, correlation time and variance are only taken into account implicitly.

Here, we present a method from a different perspective, principally based on the functional (firing) properties of the neuron under *in vivo* conditions. Our method provides an “equivalent” single compartment neuron, with very similar input-output transfer function. This reduction is obtained by focusing on three voltage moments *µ*_*v*_, *σ*_*v*_ and *τ*_*v*_, which are key to determine the transfer function. We obtain point-neuron models that have similar voltage moments, and firing characteristics as described by the transfer function.

### 4.2 Perspectives for further work

Our reduction method relies on precise voltage statistical metrics to identify the best point-neuron model with the same firing distribution. It is one of the first tools to fully reduce a morphological model into a single compartmental model. It is also an interesting tool to calculate exactly the transfer function of a neuron based on precise morphological data and build a mean field model (Zerlaut et al., 2016; El Boustani and Destexhe, 2009; Di Volo et al., 2019). The perspective of obtaining mean-field models directly from morphologically detailed models is a very promising extension of the present work.

This method is also very interesting to understand the direct effect of specific morphological properties on the firing distribution of a neuron. We can use it to understand how spatial organization or morphological properties such as active dendrites, synapse clusters, membrane area, inhomogeneous membrane parameters or channel distributions directly shape the transfer function. Our approach also accounts for the effect of signal propagation on the membrane potential which changes the shape of synaptic inputs whereas all synaptic inputs in the analytical model have the same shape. Synaptic interactions along dendrites also strongly shape membrane statistics.

Importantly, our method calculates the voltage statistics and firing rates of a detailed morphological neuron model as part of a specific network and not in an isolated artificial context such as current clamp experiments. The model is directly defined by the fitted passive properties as well as the exact synaptic currents that the neuron receives. The identified point-neuron model therefore is not supposed to have the exact same voltage dynamics close to threshold as those we would get using classic fitting procedures that rely on current injections but rather effective parameters that are guaranteed to preserve the firing rates of the neuron in its network.

This method is also very useful in comparing neurons from various animal classes, especially since neurons from a given species have very specific morphological properties. For instance, in mammals, the soma is considered as the synaptic integrator and is often directly linked to the axon, thus playing a central role in computation and spiking activity. However, in insect neurons the axon is often separated from the soma by a neurite, from which both the dendrites and axon emerge. The soma therefore seems to play no computational role at all. Our approach can be used to assess the importance of soma location on computation as well as other fundamental questions about morphological organization.

